# Prime editing primarily induces undesired outcomes in mice

**DOI:** 10.1101/2020.08.06.239723

**Authors:** Tomomi Aida, Jonathan J. Wilde, Lixin Yang, Yuanyuan Hou, Mengqi Li, Dongdong Xu, Jianbang Lin, Peimin Qi, Zhonghua Lu, Guoping Feng

## Abstract

Genome editing has transformed biomedical science, but is still unpredictable and often induces undesired outcomes. Prime editing (PE) is a promising new approach due to its proposed flexibility and ability to avoid unwanted indels. Here, we show highly efficient PE-mediated genome editing in mammalian zygotes. Utilizing chemically modified guideRNAs, PE efficiently introduced 10 targeted modifications including substitutions, deletions, and insertions across 6 genes in mouse embryos. However, we unexpectedly observed a high frequency of undesired outcomes such as large deletions and found that these occurred more often than pure intended edits across all of the edits/genes. We show that undesired outcomes result from the double-nicking PE3 strategy, but that omission of the second nick largely ablates PE function. However, sequential double-nicking with PE3b, which is only applicable to a fraction of edits, eliminated undesired outcomes. Overall, our findings demonstrate the promising potential of PE for predictable, flexible, and highly efficient *in vivo* genome editing, but highlight the need for improved variations of PE before it is ready for widespread use.

## Introduction

CRISPR/Cas-based approaches for targeted genome modification have revolutionized modern biology and hold great promise for therapeutic interventions for debilitating genetic disorders. In particular, engineered enzymes have given the gene editing field an ever-expanding set of tools with increased fidelity (Kleinstiver et al., 2016; Slaymaker et al., 2016), altered target specificities (Chatterjee et al., 2018; Hu et al., 2018; Nishimasu et al., 2018; Walton et al., 2020), the ability to directly introduce specific changes to a target genome (Gaudelli et al., 2017; Komor et al., 2016; Nishida et al., 2016), or improved genome modification capabilities (Aida et al., 2015; Charpentier et al., 2018; Gu et al., 2018; Jayavaradhan et al., 2019; Nakade et al., 2018; Rees et al., 2019). However, these options continue to suffer from low efficiencies, high indel rates, a narrow scope of editing outcomes, and/or a reliance upon restricted sets of suitable protospacer adjacent motifs (PAMs) in close proximity to the target site. A recent advance toward solving these problems was the development of the prime editing (PE) system, which enables the installation of substitutions, as well as deletions and insertions of variable sizes, into specific genomic loci with a less stringent requirement for a nearby PAM and low indel rates (Anzalone et al., 2019). Prime editing is enabled by an optimized fusion of the Moloney Murine Leukemia Virus reverse transcriptase (RT) to the H840A nickase mutant of SpyCas9 which, when paired with a guide RNA encoding the target protospacer, an RT primer binding site, the desired edit, and a short homology region (prime editing guide RNA, pegRNA), is capable of precisely writing modifications into a desired locus (Anzalone et al., 2019). The PE system was primarily developed in HEK293T cells and extensive testing at multiple loci showed high levels of installation of the intended mutations at target loci with indel rates typically less than 5%, suggesting that PE-based editing could represent a monumental step forward in the development of safe and effective gene therapies for a wide range of genetic disorders. However, additional testing in other non-primary cell lines, primary cultured mouse neurons, plants, and mice all showed lower overall rates of editing, more variable indel rates, and increased rates of insertions of pegRNA scaffold sequences (Anzalone et al., 2019; Lin et al., 2020; Liu et al., 2020), suggesting that differences in DNA repair mechanisms, cell cycle status, and chromosomal stability could have profound effects on PE outcomes. Currently, it is of great interest to understand whether PE broadly works *in vivo* for animal model production and therapeutic development (Cohen, 2019; Hampton, 2020; Ravindran, 2019; Urnov, 2020). Thus, we set out to better understand the utility of PE for editing primary cells *in vivo* and generating animal models by extensively testing its efficiency and accuracy in mouse zygotes.

## Results

### Prime editing by PE3 in mouse embryos

To begin, we designed pegRNAs to install mutations in three genes in the mouse genome: *Dnmt1, Chd2*, and *Tyr* (**Figures 1A, 1B, and S1**). We chose to start with the *Dnmt1* site because successful prime editing of this locus was previously demonstrated in cultured primary mouse cortical neurons (Anzalone et al., 2019). The *Chd2* and *Tyr* sites were chosen because we have worked extensively with these loci and have consistently observed moderate-to-high editing rates with standard SpyCas9 (Wilde et al., 2018). The initial PE study found that including a second sgRNA to induce a nick on the opposite, unedited DNA strand (>30nt from the PE site) can significantly enhance editing rates, a system referred to as PE3 (compared to PE2, which uses a single nick at the PE site) (Anzalone et al., 2019). The protospacers chosen for the second nick for *Dnmt1, Chd2*, and *Tyr* reside 53bp, 52bp, and 56bp from the on-target nicking sites, respectively (**Figures 1A, 1B, and S1**).

**Figure 1.**
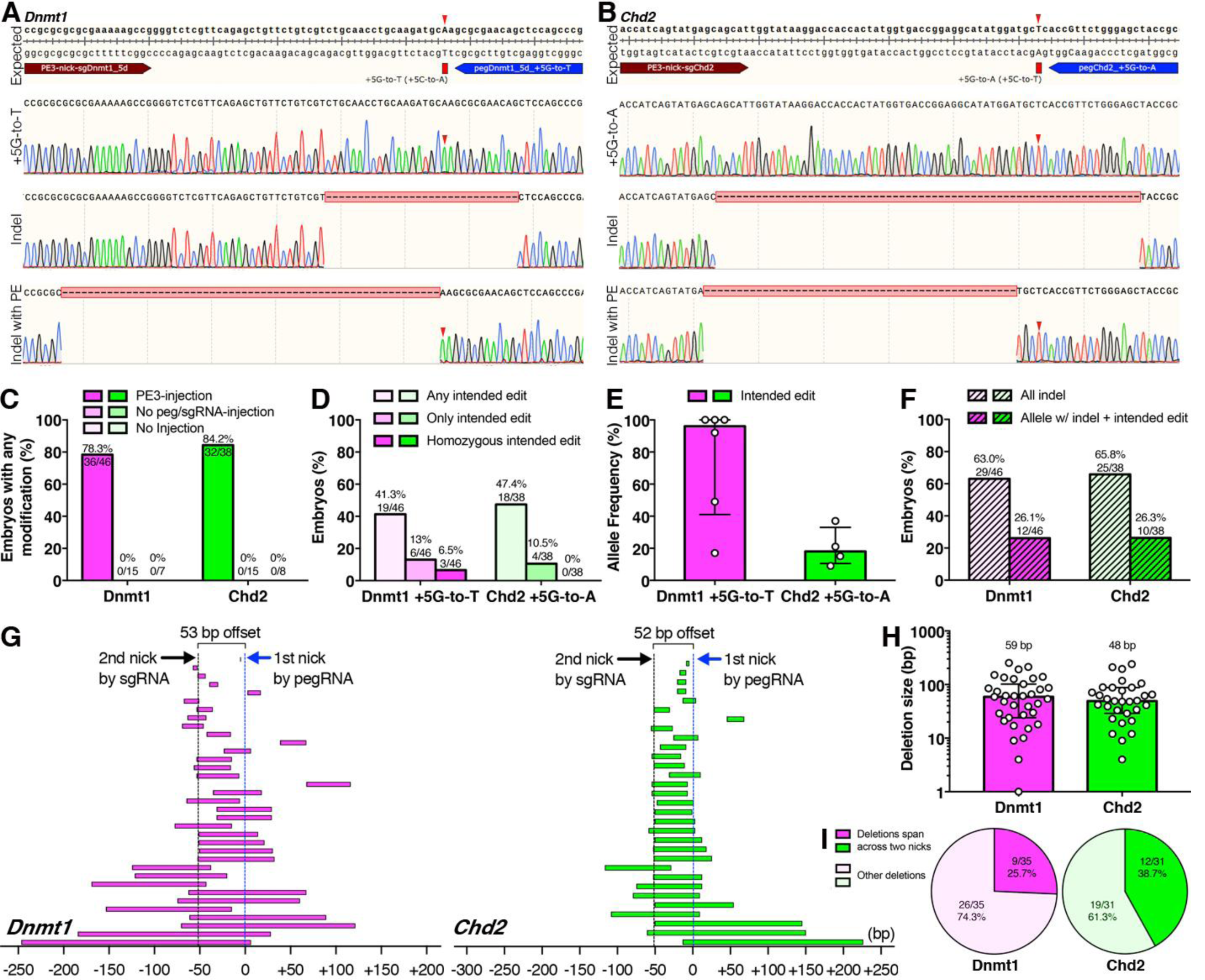
Prime editing by PE3 in mouse embryos. (**A** and **B**) Sanger sequencing chromatograms of mouse embryos injected with PE3 targeting (A) *Dnmt1* and (B) *Chd2*. Top panels: design of PE3 strategies. Mutations to be introduced are shown in upper case and indicated by red arrows. Second panels from top: cloned sequences with intended mutations (red arrows). Second panels from bottom: cloned sequences with indels (pink dashes). Bottom panels: cloned sequences with indels (pink dashes) and intended mutations (red arrows). (**C-D**) Rates of (C) any modification and (D) installation of intended edits. Alleles with intended edits were identified by TOPO cloning. (**E**) Allele frequencies of intended mutations (see **STAR methods**) shown as median ± interquartile range. (**F**) Percentage of embryos with either indels or individual alleles carrying both an indel and the intended edit. (**G**) Deletion ranges. Each bar represents a single deletion event. The pegRNA nick sites (blue dotted lines) are set as position 0. The second nicking-sgRNA target sites are indicated with black dotted lines. (**H**) Deletion sizes, shown as median ± interquartile range. (**I**) Deletion patterns.

Initial prime editing attempts using T7 *in vitro* transcribed pegRNAs and nicking sgRNAs were successful but inefficient, likely due to technical issues (**Figure S2**; see **Supplementary Discussion**). Thus, inspired by our previous success with chemically synthesized guide RNAs showing drastic improvement of homology-directed repair (HDR) efficiency in mice (Aida et al., 2015), we performed injections using chemically synthesized, chemically modified pegRNAs and nicking-sgRNAs for *Dnmt1* and *Chd2*. We observed much higher overall levels of PE3-dependent editing, with >75% of PE3-injected embryos carrying correct edits, indels, or a combination of both at the target site (**Figures 1A-1C**). We did not find any modifications of the target loci in uninjected embryos or embryos injected only with the PE2 enzyme (**Figure 1C**). Approximately 50% of PE3-injected embryos were positive for the intended edit, with 6.5% of *Dnmt1* embryos showing homozygous editing (**Figure 1D**), demonstrating the potential of prime editing for efficient *in vivo* modification. However, sequencing of individual alleles revealed significant mosaicism, with PE allele frequencies ranging from 17-100% at the *Dnmt1* locus and 9-37% at the *Chd2* locus (**Figure 1E**). Surprisingly, we found that although 41.3% (19/46) of *Dnmt1* embryos and 47.4% (18/38) of *Chd2* embryos were positive for the intended edit, only 13% (6/46) of *Dnmt1* embryos and 10.5% (4/38) of *Chd2* embryos carried the correct prime edit without the presence of indels. Unexpectedly, more than 60% of embryos harbored indels (29/46 *Dnmt1* and 25/38 *Chd2*) (**Figure 1F**). Moreover, 26.1% (12/46) of *Dnmt1* embryos and 26.3% (10/38) of *Chd2* embryos carried alleles with both the intended prime edit and an unwanted indel (**Figure 1F**). In other words, while we often observed successful installation of the intended prime edit, it was frequently nullified by the installation of unwanted genetic lesions on the same allele. In an attempt to confirm these results, we generated pups by PE3 and found similarly high indel rates (**Figure S3**). Interestingly, we observed an increase in PE rates and a decrease in indel rates at the *Dnmt1* locus (**Figures S3B** and **S3D**). However, previous studies have shown that homozygous deletion of *Dnmt1* is embryonic lethal (Li et al., 1992), suggesting that these apparent rate changes were due to the death of embryos with biallelic disruption of *Dnmt1*.

To better understand the nature of indels induced by prime editing, we analyzed the position and size of deletions with respect to the two PE3 nicks. Deletions at the *Dnmt1* locus originated from both nick sites and extended bidirectionally, with a median size of 59bp (**Figures 1G** and **1H**). Deletions at the *Chd2* locus were of similar size and also extended bidirectionally from both nick sites, with a median size of 48bp (**Figures 1G** and **1H**). Large deletions spanning the complete distance between both nicks (53bp for *Dnmt1*, 52bp for *Chd2*) made up 25.7% and 38.7% of all deletions at the *Dnmt1* and *Chd2* loci, respectively (**Figure 1I**). We also found that nearly 20% of all observed deletions were greater than 100bp in size (**Figures 1G** and **1H**).

To identify the source of these unexpected outcomes by PE3, we performed injections with engineered PE2 mRNA lacking the RT domain (Cas9 H840A nickase), pegRNA, and nicking-sgRNA, or PE2 mRNA, pegRNA lacking the RT template (RTT) and PBS (sgRNA), and nicking-sgRNA, and found frequent indel inductions consistent with PE3 injections (**Figure S4**). Furthermore, to exclude the possibility that unintentionally truncated pegRNAs lacking RTT and/or PBS during T7-IVT or chemical synthesis resulted in high frequency indels, we performed injections with DNA encoding PE2, pegRNA, and nicking-sgRNA, in which the pegRNA and nicking-sgRNA were intracellularly transcribed by the U6 promoter. Although the overall modification rates were drastically lower with DNA injections, as expected (**Figure S5A**), we observed a low level of intended editing of *Chd2* (**Figures S5B** and **S5C**) with a similar indel frequency (**Figure S5D**). Together, our data indicate that while PE3 is capable of installing intended prime edits at high rates in mouse zygotes, the primary outcome is large, unwanted DNA lesions that result from the double nicking strategy of PE3.

### Expanded analysis of prime editing at additional loci

Next, we asked whether the outcomes of PE3-mediated genome editing can be generalized across loci in mice by testing 9 additional prime editing conditions for 6 edits across 4 new loci (**Figure 2**). After attempting to map all pegRNAs and nicking-sgRNAs used by Anzalone *et al* onto the mouse genome, we found that *Rnf2*, which accounts for 22% of all editing reported in that study, is a conserved target between human and mouse. At this locus, we attempted 3 different edits: a single base substitution (+1C-to-G), a 3bp insertion (+1GTAins), and a 3bp deletion (+3-5GAGdel) in combination with 2 different nicking-sgRNAs (+41 and +67) from the previous report (Anzalone et al., 2019). We also tested edits in 3 genes that we previously targeted by HDR: a two base substitution (+45TC-to-AA) in *Tyr* as described above (Wilde et al., 2018), a 6bp insertion (+1GAATTCins, insertion of EcoRI site) in *Actb* (Aida et al., 2015, 2016), and a single base substitution (+2A-to-C) in *Col12a1* (Aida et al., 2016; Quadros et al., 2017).

**Figure 2.**
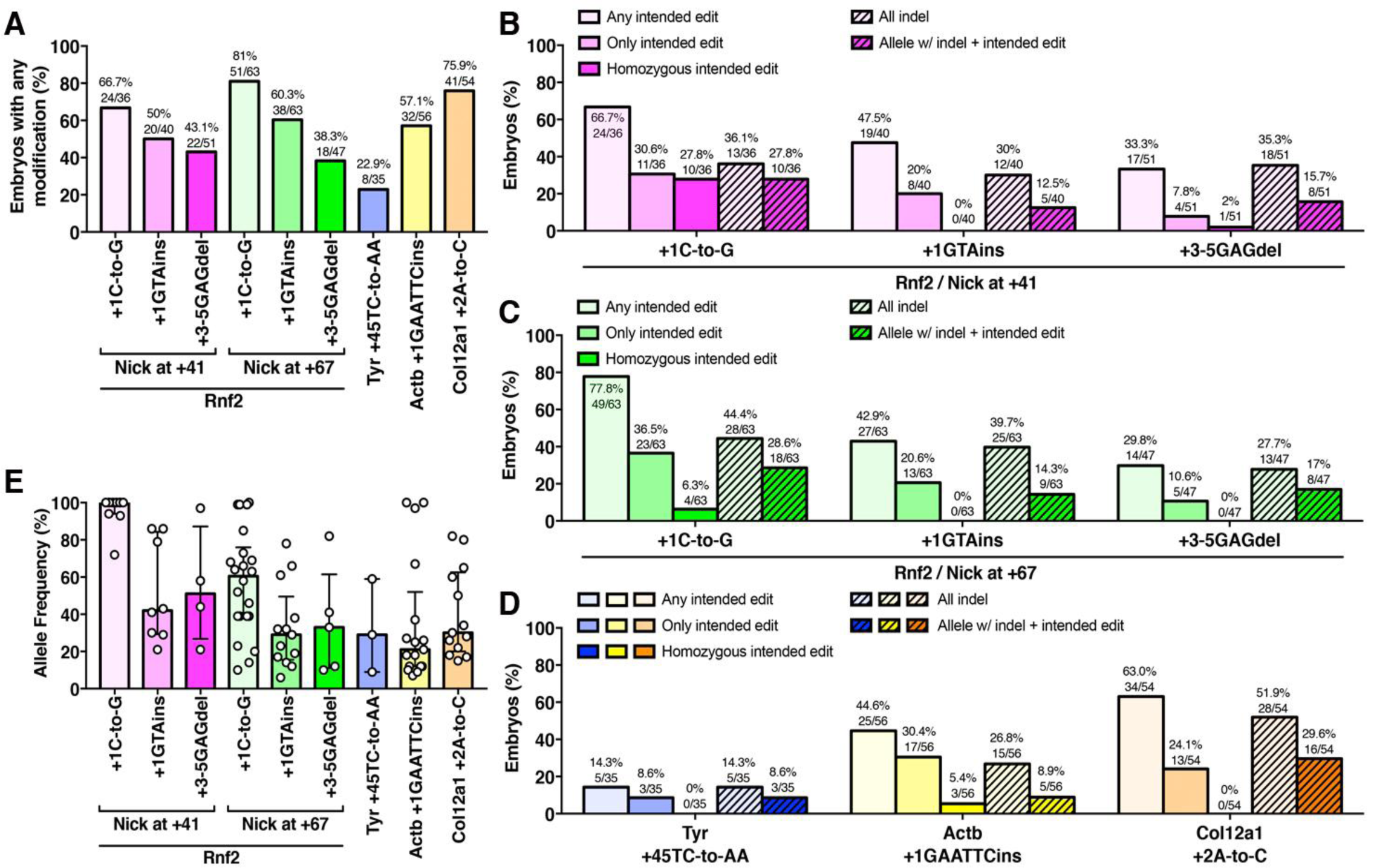
Variable prime editing outcomes by PE3 across genes in mouse embryos. (**A-E**) Rates of (A) any modification and (B-D) installation of intended edits, indels, and individual alleles carrying both an indel and the intended edit. Alleles with intended edits were identified by TOPO cloning. (**E**) Allele frequencies of intended mutations (see **STAR methods**) shown as median ± interquartile range.

We observed moderate-to-high overall levels of PE3-dependent editing for all 9 conditions, creating 6 edits across 4 loci including the previously uneditable *Tyr* (**Figures 2A** and **S2**) by using chemically synthesized pegRNAs. Approximately 70% (24/36 and 49/63) of PE3-injected embryos were positive for the +1C-to-G substitution in *Rnf2* by either nicking-sgRNA at +41 or +67, respectively, with a higher homozygous rate by nicking-sgRNA at position +41 (27.8%, 10/36 by nicking-sgRNA at +41 compared to 6.3%, 4/63 nicking-sgRNA at +67: p<0.01), demonstrating highly efficient PE for +1C-to-G installation in *Rnf2* (**Figures 2B** and **2C**). We also observed high median allele frequencies of more than 90% by nicking-sgRNA at +41 and more than 50% by nicking-sgRNA at +67 (**Figure 2E**). However, we also found frequent indels in both cases, (36.1%, 13/36 by nicking-sgRNA at +41 and 44.4%, 28/63 by nicking-sgRNA at +67) with indel rates higher than those for only the intended edit (30.6%, 11/36 by nicking-sgRNA at +41 and 36.5%, 23/63 by nicking-sgRNA at +67) (**Figures 2B** and **2C**).

We observed similar outcomes for the +1GTAins and +3-5GAGdel edits in *Rnf2* by either nicking-sgRNA at +41 or +67, although the overall modification rates and rates of intended edits were lower (**Figures 2B** and **2C**), suggesting that PE efficiency may be higher for single nucleotide substitutions than insertions or deletions. Importantly, we once again observed higher rates of indels than of correct edits, demonstrating that indel formation is independent from the type of mutation being installed (**Figures 2B** and **2C**). Furthermore, we found similar results for 3 additional edits at 3 independent loci: the total modification rates and rates of intended edits were 22.9% (8/35) of total modification and 14.3% (5/35) of intended edits for +45TC-to-AA in *Tyr*, 57.1% (32/56) of total modification and 44.6% (25/56) of intended edits for +1GAATTCins in *Atcb*, and 75.9% (41/54) of total modification and 63% (34/54) of intended edits for +2A-to-C in *Col12a1*, respectively (**Figures 2A, 2D** and **2E**). Consistent with *Rnf2*, we observed equivalent or higher rates of indels than the rates of pure intended edits for all edits/loci (14.3% (5/35) of all indel and 8.6% (3/35) of pure intended edits for +45TC-toAA in *Tyr*, 26.8% (15/56) of all indel and 30.4% (17/56) of pure intended edits for +1GAATTCins in *Atcb*, and 51.9% (28/54) of all indel and 24.1% (13/54) of pure intended edits for +2A-to-C in *Col12a1*, respectively) (**Figure 2D**), demonstrating that indel formation is a predominant editing outcome by PE3 in mouse zygotes that is independent of locus and intended mutation type. Altogether, we demonstrate highly efficient genome editing in mouse embryos by PE3 using chemically synthesized pegRNAs for 11 editing strategies across 6 genes, including 7 previously characterized editing strategies in 2 genes, but unexpectedly revealed that indel formation occurs as often or more often than pure prime editing at all edits/loci tested.

### Prime editing by PE2 in mouse embryos

While the double nicking strategy employed by PE3 does not directly induce double-strand breaks, a clear understanding of how these nicks are processed and repaired is lacking and our data suggest that the double nicking may be responsible for the large indels we observed (**Figure 1G**). We tried to understand the role of the second nick in undesired indel formation in detail by titrating nicking-sgRNA levels *in vivo*, since the initial *in vitro* prime editing experiments used a 3:1 ratio of pegRNA to nicking-sgRNA for transfections (Anzalone et al., 2019) and we used an ∼1:1 ratio (50 ng/μl each) *in vivo*. We performed injections using 50, 17, 6, and 2 ng/μl nicking-sgRNA while maintaining a pegRNA concentration of 50 ng/μl for the *Dnmt1* and *Chd2* edits in order to test 1:1, 3:1, 8:1, and 25:1 ratios of pegRNA to nicking-sgRNA (**Figures 3A-D**). As nicking-sgRNA concentrations decreased, we observed increased rates of embryos with pure intended edits and conversely decreased rates of embryos with indels (**Figures 3B** and **3D**). These results strongly suggest that the rates of intended edits and undesired indels are dependent upon nicking-sgRNA levels. However, even with the lowest concentration (2 ng/μl) of nicking-sgRNA we still observed indel rates higher than 45% at both loci (**Figure 3D**), suggesting that indel formation may be an inherent feature of PE3 that cannot be overcome by simply decreasing the concentration of nicking-sgRNA.

**Figure 3.**
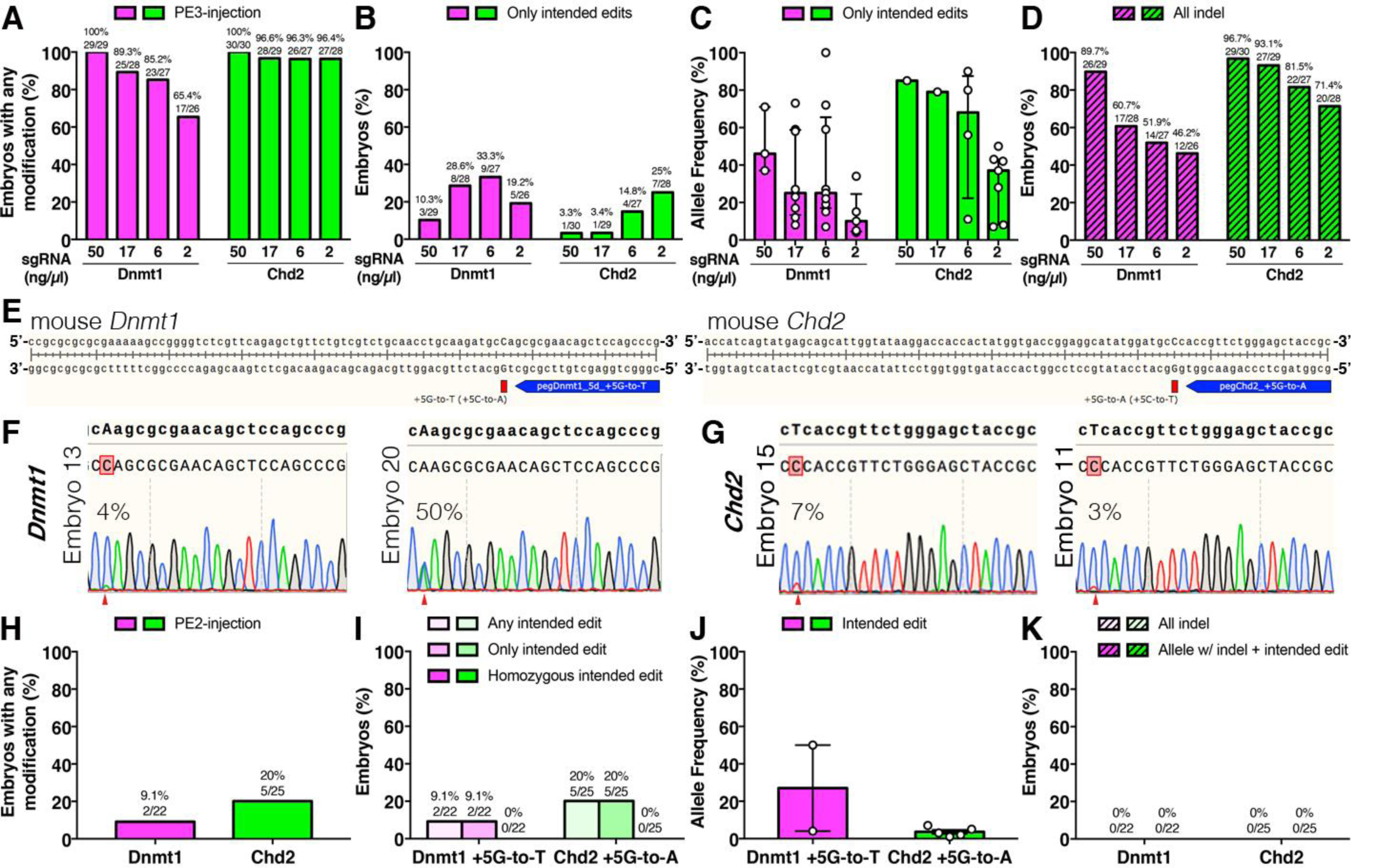
Prime editing by PE2 in mouse embryos. (**A-D**) Titration of nicking-sgRNA. Rates of (A) any modification, (B) installation of only intended edits, and (D) indels. (C) Allele frequencies of intended mutations (see **STAR methods**) shown as median ± interquartile range. (**E**) Design of PE2 strategies. (**F** and **G**) Sanger sequencing chromatograms of mouse embryos injected with PE2 targeting (F) *Dnmt1* and (G) *Chd2*. Representative embryos are shown. Top text sequences: intended editing outcomes with the targeted mutations (upper case). Bottom text and chromatogram: Sanger results. Expected mutations are shown by red arrows. The “C”s with pink highlights indicate dominant wild-type “C” peaks at target sites. Numbers indicate allele frequencies of the intended mutations (see **STAR methods**). (**H** and **I**) Rates of (H) any modification and (I) installation of intended edits. Alleles with intended edits were identified by TOPO cloning. (**J**) Frequency of alleles with intended edit are shown as median ± interquartile range. (**K**) Indel rates.

Although initial experiments with PE showed a general increase in PE efficiency when including a second nick, multiple prime edits were also efficiently introduced without it in a nearly indel-free manner (Anzalone et al., 2019). We therefore targeted the *Dnmt1* and *Chd2* loci in mouse zygotes with this so-called PE2 strategy, which does not utilize the second nick, to assess whether it could induce precise editing without indels. At the *Dnmt1* locus we observed editing in just 2/22 embryos and at the *Chd2* locus there was editing in 5/25 embryos (**Figures 3E-H**). However, in 6/7 edited embryos the frequency of the prime edited allele was less than 10%, while one embryo was heterozygous for the intended edit (**Figures 3I** and **3J**), indicating a substantial decrease in efficiency compared to PE3. In support of the hypothesis that the second nick included in PE3 stimulates indel formation, we did not observe any indels in the injected embryos (**Figure 3K**). These observations are in agreement with previous work that has described indels resulting from double nicking (Mali et al., 2013; Ran et al., 2013). Taken together, these data show that while a second nick is required for efficient induction of prime editing, it is also the primary driver of PE3-stimulated indel formation.

### Comparison with Homology-Directed Repair by wild-type SpyCas9-RNP and ssODN donor in mouse embryos

Despite the frequent induction of indels by PE3, its efficiency of installing the intended edit led us to investigate whether prime editing still has advantages over the traditional ribonucleoprotein (RNP)-based HDR method (Aida et al., 2015) commonly used for knock-in mouse production (Abe et al., 2020; Quadros et al., 2017). We designed single-stranded oligodeoxynucleotide (ssODN) donors for the same *Dnmt1* and *Chd2* edits (Wilde et al., 2018) made with PE3 and co-injected them with RNPs consisting of SpyCas9 protein and sgRNAs targeting the same on-target nick sites used for prime editing (**Figure 4A**). We observed knock-in of the intended allele, as well as indels, at both loci using this method and modification of the target site was >80% for both loci (**Figures 4B-D**). At the *Dnmt1* locus 48% (12/25) of embryos contained the intended edit, while 73.9% (17/23) contained the intended edit at the *Chd2* locus, consistent with our previous findings (Wilde et al., 2018). Of these, approximately 50% contained the intended edit without indels and nearly all of these embryos (13/14) were homozygous for the correct knock-in (**Figure 4E**), with overall HDR allele frequencies ranging from 97-100% at the *Dnmt1* locus and 92-100% at the *Chd2* locus (**Figure 4F**). We observed indels in 15/25 *Dnmt1* embryos and 11/23 *Chd2* embryos but in contrast to PE3, only 5/48 occurred on the allele carrying the intended edit (**Figure 4G**). Because we frequently observed large deletions as a result of prime editing, we analyzed the size and position of indels resulting from HDR injections and found significant differences (**Figures 4H-J** and **S6**). While the median PE3-generated deletion sizes for *Dnmt1* and *Chd2* were 59bp and 48bp, respectively (**Figure 1H**), the median sizes at the same sites after DSB induction by SpyCas9 were only 15bp and 8bp (**Figures 4I** and **S6**). Additionally, we observed a shift in the directionality of indels between PE3 and HDR (**Figure 4H**). While PE3-based editing caused indels extending primarily in the direction of the second nick (**Figure 1G**), indels observed in our HDR experiments primarily extended in the opposite direction of the putative nick site at the *Dnmt1* locus and did not show a directional preference at the *Chd2* locus (**Figure 4H**). While a second nick was not utilized in the HDR experiments, the size and patterns of observed indels further support the conclusion that the second nick required for efficient prime editing is responsible for the frequency, size, and directionality of PE3-associated indels. Together, these data indicate that prime editing in its current form does not provide a significant advantage over traditional HDR methods in mouse embryos.

**Figure 4.**
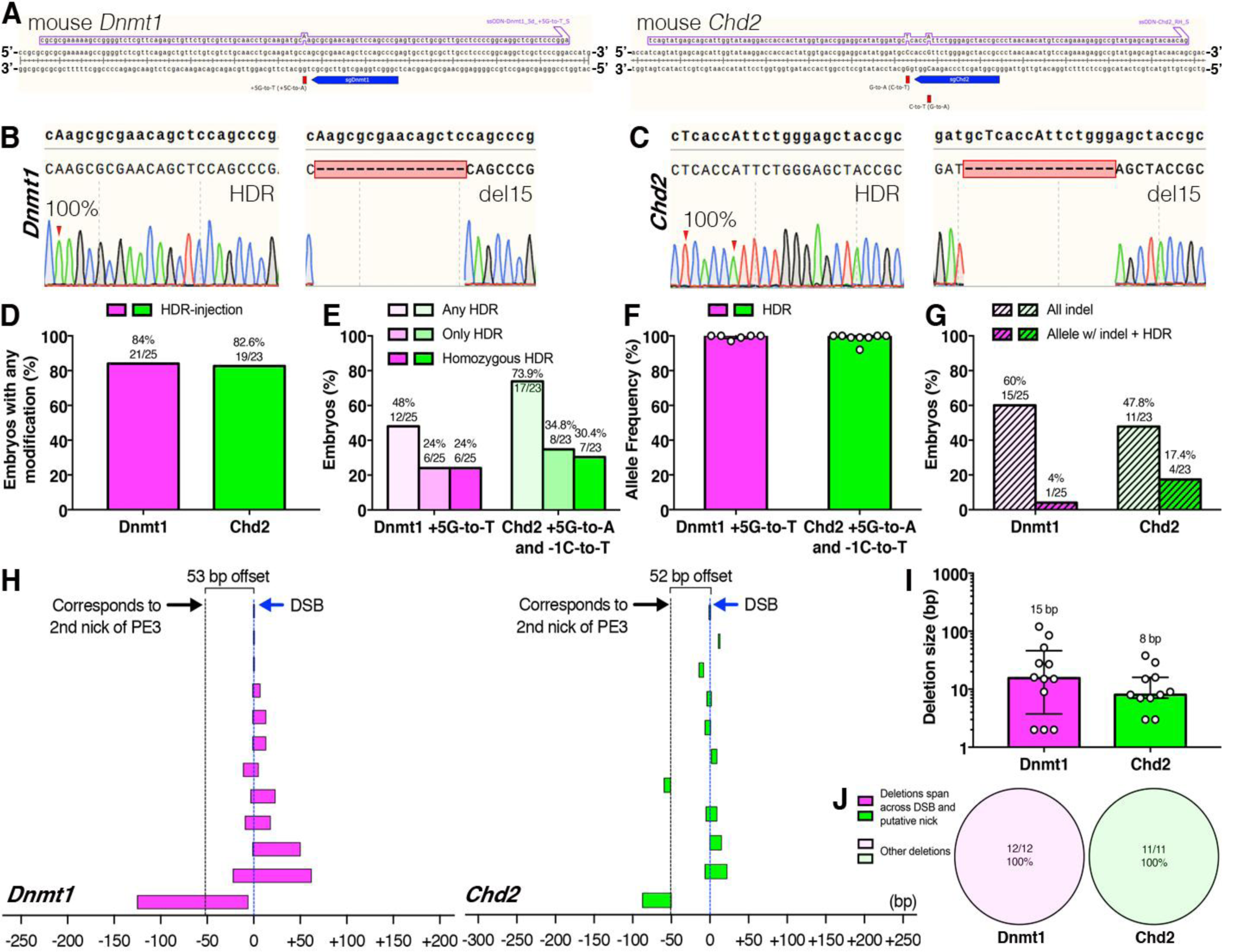
Homology-Directed Repair (HDR) by wild-type SpyCas9-RNP and ssODN donor in mouse embryos. (**A**) Design of HDR strategies. Top sequences: ssODN donors. (**B** and **C**) Sanger sequencing chromatograms of mouse embryos injected with CRISPR HDR reagents targeting (B) *Dnmt1* and (C) *Chd2*. Sanger chromatograms for HDR (bulk) and indels (cloned) from representative embryos are shown. Top text sequences: expected editing outcomes with the targeted mutations (upper case). Bottom: Sanger traces. Expected mutations are shown by red arrows. Numbers indicate allele frequencies of the intended mutations (see **STAR methods**). Pink dashes indicate indels. (**D-E**) Rates of (D) any modification and (E) HDR. (**F**) Allele frequencies of the intended mutations as median ± interquartile range. (**G**) Percentage of embryos with either indels or individual alleles carrying both an indel and the intended edit. (**H**) Deletion ranges. Each bar represents a single deletion event. The ‘0’ positions are set at the DNA double-strand break (DSB) sites (blue dotted lines). The putative second nick sites for sgRNAs used in PE3 are also indicated with black dotted lines for comparison. (**I**) Deletion sizes with median ± interquartile range shown. (**J**) Deletion patterns.

### Prime editing by PE3b in mouse embryos

In an attempt to maintain high levels of prime editing while reducing indels, a modified version of prime editing called PE3b, which relies upon sequential double nicking mediated by installation of the intended mutation on the target strand, which when edited destroys the original nicking site and creates a new nicking site on the opposite strand, was developed (Anzalone et al., 2019). We tested this method at the *Dnmt1* and *Chd2* loci in mouse embryos to determine whether it provides a solution for PE-mediated indel formation. Due to the limited flexibility of nicking-sgRNA design with PE3b, which can only be used when installation of the intended mutation generates a new PAM for the nicking-sgRNA on the unedited strand and destroys the PAM or protospacer associated with the pegRNA, we were not able to use this strategy to install the mutations previously used in this study. Therefore, we introduced new mutations that destroy the pegRNA PAM sites and create novel PAM sites for the PE3b nicking-sgRNA at the same loci (**Figure 5A**). Target site modification rates were slightly altered compared to PE3, with 91.7% (55/60) and 47.5% (29/61) embryos showing signs of editing at the *Dnmt1* and *Chd2* loci, respectively (**Figures 5B-D**). However, of the embryos showing modification of the target site, nearly 100% were positive for the intended edit (**Figure 5E**). At the *Dnmt1* locus we observed variable mosaicism, indicated by allele frequencies ranging from 11% to 100% (median 54%, **Figure 5F**). Editing rates also varied at the *Chd2* locus, indicated by allele frequencies ranging from 5% to 66% (median 16%, **Figure 5F**). Nonetheless, we observed indels in only 2/60 *Dnmt1* embryos and 2/61 *Chd2* embryos, demonstrating the usefulness of PE3b for eliminating indels associated with PE3 (**Figure 5G**). Therefore, we conclude that PE3b is a viable option for reducing or eliminating indels associated with prime editing but note that its design restrictions, which limit the number of mutations for which it can be applied, necessitate additional advances to improve prime editing for applications that require high efficiency and allele frequency (Aida and Feng, 2020; Li et al., 2020; Zhang et al., 2020; Zhou et al., 2019).

**Figure 5.**
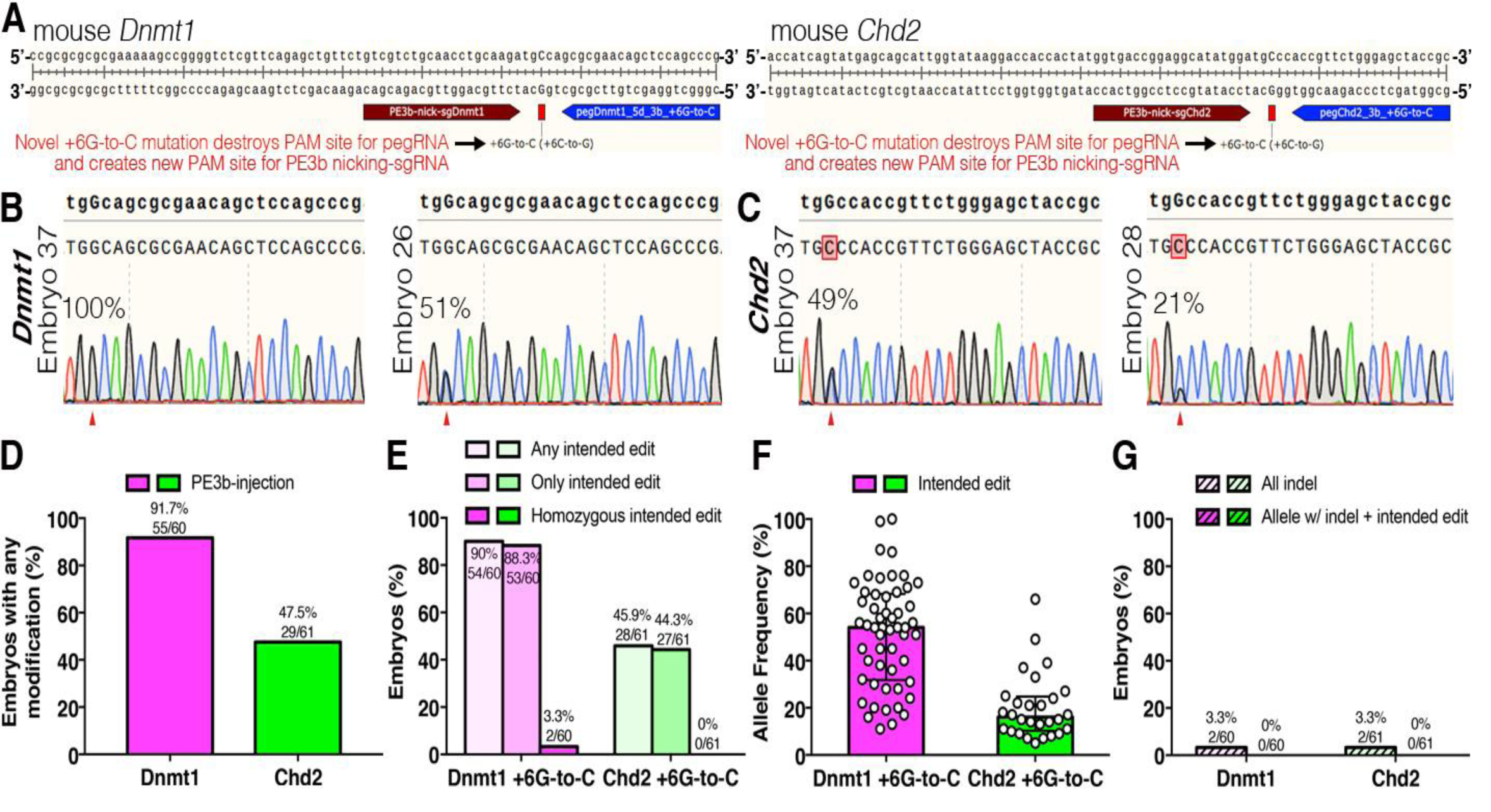
Prime editing by PE3b in mouse embryos. (**A**) Design of PE3b strategies. Successful prime edits destroy the pegRNA PAM sites and introduce new PAM sites for the nicking-sgRNA. Note that PE3b introduces different mutations than those introduced by PE3, PE2, or HDR. (**B** and **C**) Sanger sequencing chromatograms of mouse embryos injected with PE3b targeting (B) *Dnmt1* and (C) *Chd2*. Representative embryos are shown. Top text sequences: intended editing outcomes with the targeted mutations (upper case). Bottom text and chromatogram: Sanger results. Expected mutations are shown by red arrows. The “C”s with pink highlights indicate dominant wild-type “C” peaks at target sites. Numbers indicate allele frequencies of the intended mutations (see **STAR methods**). (**D** and **E**) Rates of (D) any modification and (E) installation of intended edits. (**F**) Frequency of alleles with intended edit are shown as median ± interquartile range. (**G**) Indel rates.

## Discussion

While development of the prime editing platform was an exciting step forward for gene editing, our study shows that there are significant differences between editing outcomes in cultured cells and primary mouse embryos. Although the exact reason for these discrepancies is not completely clear, our data clearly demonstrate that the second nick required for efficient editing with PE3 is the primary driver of indel formation *in vivo*. While our work does demonstrate PE3’s ability to efficiently install targeted mutations in mouse embryos, high indel rates and frequent unwanted mutations on the prime edited allele largely negate its utility for animal model generation or gene therapy in its current form.

Nicks are primarily repaired through the high-fidelity base excision repair pathway and are thus non-mutagenic, although they can be converted to DSBs during DNA replication (Dianov and Hübscher, 2013). However, this is unlikely to occur frequently with prime editing since we did not observe any indels in mouse embryos edited by PE2, which induces a single nick. The PE3 strategy that requires induction of nicks on both DNA strands has also been utilized to induce DSBs and subsequent indels by zinc finger nucleases (Miller et al., 2007), transcription-activator-like effector nucleases (Miller et al., 2011), and CRISPR Cas9-D10A nickases (Mali et al., 2013; Ran et al., 2013). Therefore, it is not surprising that induction of paired nicks in the PE3 context results in DSBs and subsequent indels. We show that both titration of nicking-sgRNAs and sequential, mutually exclusive nicking with PE3b reduce indels while preserving some prime editing function, strongly suggesting that induction of paired nicks at the same time results in indels. We and others have found additional evidence of dual nicking-induced errors in the frequent observation of intended edits with indels on the same allele (Lin et al., 2020). This potentially explains at least one mutagenic DNA repair mechanism by PE3: 1) the PE2 enzyme efficiently reverse transcribes the pegRNA at the first nick, 2) integration of the 3’ flap containing the intended edit into the genome as predicted, 3) loss of genetic information from the flanking strands to be ligated with the integrated 3’ flap, potentially due to 5’ end resection between double nicks, which inserts indels between the integrated 3’ flap and the second nick made by the nicking sgRNA. Since the 5’ flap (unedited DNA) is thought to be removed by 5’ exonucleases such as EXOI (Anzalone et al., 2019; Keijzers et al., 2015), it’s possible that EXO1 also stimulates longer 5’ end resection and microhomology mediated end-joining (Aida et al., 2016).

Our work and the published prime editing studies discussed in this manuscript describe clear discrepancies between editing outcomes in cultured cells and *in vivo*. We highlight these discrepancies using 7 validated pegRNA/nicking-sgRNA combinations previously shown to have low indel rates in HEK293T cells, suggesting that there is much to be learned about the mechanism of prime editing. Our results show the formation of undesired DSBs and subsequent indels that are mediated by the double nicking strategy of PE3, which is critically impacted by dose, sgRNA activity, and timing of the second nick. Compared to cultured cells where PE components are heterogeneously transfected into each cell, microinjection delivers the components into mouse zygotes in a relatively homogeneous and simultaneous manner, potentially explaining the high indel rates in our study. However, the least understood aspect of prime editing is the repair mechanism employed at the target site, which is likely to have significant impacts on prime editing outcomes. While it is hypothesized that this is mediated by canonical flap repair pathways, particularly those relying upon flap endonuclease 1 (FEN1), these hypotheses have not been directly tested and it’s likely that the activity and availability of many DNA repair proteins affects the outcome of prime editing across loci and cell types. Previous studies have shown the *FEN1* expression is tightly linked to the cell cycle (Warbrick et al., 1998) and that tissues across the body show highly variable expression of both *FEN1* mRNA and FEN1 protein, with muscle tissues showing no detectable FEN1 expression (Uhlén et al., 2015, Human Protein Atlas available from http://www.proteinatlas.org). Furthermore, data from GTEx (Release V6) indicate that expression of *FEN1* is abnormally high in transformed cells and cultured fibroblasts compared to primary tissue, suggesting additional sources for the discrepancy between data in HEK293T cells and *in vivo*. However, RNA-seq data indicate that *FEN1* expression is similar between HEK293 cells and U2-OS cells (Uhlén et al., 2015, Human Protein Atlas available from http://www.proteinatlas.org), which show significantly different PE rates when comparing the same sites (Anzalone et al., 2019), suggesting that additional factors contribute to the efficacy of prime editing. It is therefore critical to understand more about the DNA repair mechanisms engaged by cells to repair prime edited loci, as this likely holds the key to improving both the efficiency and accuracy of prime editing, which is critical if the technology is to be used for therapeutic purposes. Overall, we find that prime editing is capable of efficiently installing a wide range of desired mutations in mammals, but that this is often accompanied by unwanted genetic lesions that negate its success. Thus, while this technology has the potential to correct a wide range of disease-causing mutations and could be broadly useful for gene therapy, additional advances in both the technology itself and our understanding of the PE mechanism are required before this method is truly ready for prime time.

## Acknowledgements

We would like to thank Kevin Holden, John Walker, Anastasia Kadina, and Synthego for providing the synthetic pegRNAs and sgRNAs used in this study. We are grateful for assistance from the staff of MIT’s Division of Comparative Medicine. Additionally, we would like to thank Martin Wienisch, Minqing Jiang, and the rest of the Feng Lab and Lu Lab for helpful discussions and technical support regarding this work. J.J.W. is supported by a J. Douglas Tan postdoctoral fellowship, established by K. Lisa. Yang. This work was funded by the James and Patricia Poitras Center for Psychiatric Disorders Research at MIT (G.F.), the Hock E. Tan and K. Lisa Yang Center for Autism Research (G.F.)., Ministry of Science and Technology of China grant JCYJ20180504165804015 (Z.L.), Chinese Academy of Sciences International Partnership Program 172644KYSB20170004.

## Author contributions

T.A. and J.J.W. conceived and designed the experiments; T.A., L.Y., M.L., D.X., J.L., and P.Q. prepared and performed injections; T.A., L.Y., Y.H., and J.L. performed embryo and mouse tissue collections and genotyping; T.A., J.J.W., L.Y., and Z.L. analyzed data; T.A. and J.J.W. prepared the manuscript; Z.L. and G.F. supervised the project.

## Data and materials availability

Raw data from all experiments is available upon request.

## STAR Methods

### RESOURCE AVAILABILITY

#### Lead Contact

Further information and requests for resources and reagents should be directed to the Lead Contact, Guoping Feng.

#### Materials Availability

This study did not generate novel or unique materials.

#### Data and Code Availability

Raw data for the figures in this manuscript can be obtained from the Lead Contact or corresponding authors upon request.

### EXPERIMENTAL MODEL AND SUBJECT DETAILS

Mouse work was carried out in two separate animal facilities at Massachusetts Institute of Technology (MIT) in the USA, and Shenzhen Institutes of Advanced Technology (SIAT), Chinese Academy of Sciences in China. All mouse work at MIT was performed with the supervision of the Massachusetts Institute for Technology Division of Comparative Medicine (DCM) under protocol 0519-027-22, which was approved by the Committee for Animal Care (CAC). All procedures were in accordance with the guidelines set forth by the Guide for Care and Use of Laboratory Animals, National Research Council, 1996 (institutional animal assurance no. A-3125-01). All procedures at SIAT were approved by the Institutional Animal Care and Use Committee (IACUC) at the Shenzhen Institutes of Advanced Technology, Chinese Academy of Sciences.

### METHOD DETAILS

#### RNA Synthesis

All pegRNAs and sgRNAs were chemically synthesized by Synthego and Integrated DNA Technologies with chemical modifications of 2’-O-methyl analogs and 3’ phosphorothioate internucleotide linkages at the first three 5’ and 3’ terminal RNA residues (see **Table S1** for sequences).

For pegRNA synthesis by *T7-in vitro* transcription, pU6-pegRNA-GG-acceptor plasmid (Addgene #132777) was digested with BsaI-HFv2 (New England Biolabs) and the vector backbone was gel purified using the Zymoclean Gel DNA Recovery Kit (Zymo Research). T7 promoter-conjugated protospacer, 5’-phosphorylated tracrRNA scaffold, and 3’ extension oligos (Integrated DNA Technologies, see **Table S2** for sequences) were cloned into the vector by Golden Gate assembly using Quick Ligation Kit (New England Biolabs) as described previously (Anzalone et al., 2019). Cloned pU6-T7-pegRNA plasmids were digested by HindIII-HF (New England Biolabs), gel purified, and used for T7-mediated *in vitro* transcription with MEGAshortscript T7 Transcription Kit (Thermo Fisher Scientific) according to manufacturer’s instructions. pegRNAs were purified via MEGAclear kit (Thermo Fisher Scientific), aliquoted to a single-use volume, and stored at −80°C until use.

For sgRNA synthesis by *T7-in vitro* transcription, annealed protospacer oligos (see **Table S2** for sequences) were cloned into pX458 plasmid (Addgene #48138) digested with BbsI-HF (New England Biolabs). sgRNA cassettes were PCR amplified using T7-promoter conjugated primers using Herculase II Fusion DNA Polymerases (Agilent) as described previously (Aida et al., 2015; Wang et al., 2013). T7-nicking-sg*Tyr* amplicon was directly PCR amplified from empty pX458 using T7-promoter conjugated primers (see **Table S2** for sequences). sgRNA PCR amplicons were purified using DNA Clean & Concentrator-5 (Zymo Research), and used for T7-mediated *in vitro* transcription and purified as described above.

For PE2 mRNA synthesis, pCMV-PE2 plasmid (Addgene #132775) was linearized with PmeI (New England Biolabs), purified with DNA Clean & Concentrator-5 (Zymo Research), and used for T7-mediated *in vitro* transcription with the T7 mScript Standard mRNA Production System (Cellscript) according to manufacturer’s instructions. PE2 mRNA was purified via MEGAclear kit (Thermo Fisher Scientific), aliquoted to a single-use volume, and stored at −80°C. For PE2 lacking RT domain (Cas9 H840A nickase) mRNA synthesis, pCMV-PE2 plasmid was digested with BamHI-HF and EcoRI-HF (New England Biolabs) to remove RT domain, re-conjugated by using bridging oligos (see **Table S2** for sequences) and NEBuilder HiFi DNA Assembly Master Mix (New England Biolabs), and used for T7-mediated *in vitro* transcription as described above.

The concentrations and qualities of PE2 mRNA, *in vitro* transcribed pegRNA and sgRNA, and chemically synthesized pegRNA and sgRNA were checked by NanoDrop (Thermo Fisher Scientific), Bioanalyzer (Agilent), and 1% agarose gel electrophoresis.

#### in vitro Fertilization

*in vitro* fertilization (IVF) was performed using FERTIUP Mouse Preincubation Medium and CARD MEDIA (Kyudo Company) according to the manufacturer’s instructions. Non-virgin C57BL/6N males at MIT and C57BL/6J males at SIAT that were >8 weeks old were used as sperm donors. Following IVF, zygotes were cultured for 8 hours and then subjected to microinjection.

#### Injection Mixtures

Injection mixtures were prepared as described previously (Aida et al., 2015). For prime editing by RNA, nuclease-free water (Thermo Fisher Scientific), Tris-HCl, pH 7.39 (final concentration 10 mM, Thermo Fisher Scientific), PE2 mRNA (final concentration 100 ng/μL), pegRNA (final concentration 50 ng/μL), and/or nicking-sgRNA for PE3 (final concentration 50 ng/μL) were mixed. For titration of nicking-sgRNA, 50, 17, 6 and 2 ng/μl were used. For prime editing by DNA, nuclease-free water (Thermo Fisher Scientific), Tris-HCl, pH 7.39 (final concentration 10 mM, Thermo Fisher Scientific), pCMV-PE2 (final concentration 5 ng/μL), pU6-pegRNA (final concentration 1.3 ng/μL), and U6-sgRNA PCR amplicon for PE3 (final concentration 0.2 ng/μL) were mixed. For HDR using Cas9-RNP complexes, nuclease-free water (Thermo Fisher Scientific), Tris-HCl, pH 7.39 (final concentration 10 mM, Thermo Fisher Scientific), EnGen Cas9 NLS, *S. pyogenes* (New England Biolabs, final concentration 60 ng/μL), and sgRNA (final concentration 0.61 μM) were mixed and incubated at 37°C for 15 minutes, then mixed with ssODN (final concentration 100 ng/μL, Integrated DNA Technologies, see **Table S2** for sequences). RNP injection mixtures were briefly heated to 37°C prior to injection.

#### Microinjection

Microinjections were performed as described previously (Aida et al., 2015; Wilde et al., 2018). Prime editing mixtures by RNA were injected into the cytoplasm, and prime editing mixtures by DNA and Cas9-RNP complexes for HDR were injected into the male pronucleus. All microinjections were performed using a micromanipulator (Narishige), equipped on a Nikon Eclipse TE2000-S microscope at MIT or Nikon Ti2-U microscope at SIAT and an Eppendorf 5242 microinjector at MIT or Eppendorf FemtoJet 4i at SIAT. Individual zygotes were injected using an “automatic” injection mode set according to needle size and adjusted for clear increase in pronuclear volume. Following injections, zygotes were cultured in KSOM-AA under mineral oil at 37°C and 5% CO2 until collection at morula or blastocyst stage for embryonic genotyping. For generation of pups, embryos were transferred into pseudopregnant CD-1 females (Charles River Laboratories, Strain Code 022) 24-hours post-injection and allowed to develop naturally until natural birth.

#### Embryo Collection and Genomic DNA Extraction

Embryo collection and genomic DNA extraction were performed as described previously (Wilde et al., 2018). Briefly, embryos were collected at morula-to-blastocyst stage into 4-6μL of nuclease-free water (Thermo Fisher Scientific). After collection, an equivalent volume of of 2X embryo lysis buffer was added to each sample (final concentrations: 125μg/mL proteinase K (Millipore Sigma), 100mM Tris-HCl (pH 8.0, Thermo Fisher Scientific), 100mM KCl (Thermo Fisher Scientific), 0.02% gelatin (Thermo Fisher Scientific), 0.45% Tween-20 (Millipore Sigma), 60μg/mL yeast tRNA (Thermo Fisher Scientific)) and embryos were lysed for 1 hour at 56°C. Proteinase K was inactivated via incubation at 95°C for 10 minutes and DNA was stored at −20°C until use. For newborn mice, tail tissue (∼0.5 cm length) was collected from individual animals, then lysed and subjected to genomic DNA purification using the NucleoSpin Tissue kit (Macherey Nagel) according to manufacturer’s instructions.

#### PCR

PCR was performed as described previously (Wilde et al., 2018). For embryonic genotyping, two-round PCR with a nested approach was used. The first round of PCR was performed using 4-5μL of embryonic DNA as an input. For all embryo genotyping, reaction mixtures were set up in a UV-sterilized laminar flow hood using sterile reagents and filter tips to avoid contamination. All PCR was performed using nuclease-free water (Thermo Fisher Scientific), Biolase DNA Polymerase (Bioline), 10xNH4 Reaction Buffer (Bioline), 50mM MgCl2 (Bioline), and fresh aliquots of dNTPs (New England Biolabs). The second round nested PCR was then performed using 2-3μL of the product from the first round in a of PCR reaction as input. All the primers used in this study are listed in **Table S3**. For newborn mouse genotyping, a single round of PCR was performed using 2 μL of purified genomic DNA. PCR amplicons were checked on 2% agarose gel before sequencing.

#### TOPO Cloning

PCR amplicons were used as input for TA cloning using the TOPO TA Cloning Kit (Thermo Fisher Scientific) at MIT or pEASY-T1 Simple Cloning Kit (TransGen Biotech) at SIAT according to manufacturer’s instructions. Bacterial colonies were screened by blue-white selection followed by colony PCR or plasmid purification, and subjected for Sanger sequencing.

#### Sanger Sequencing

For all sequencing reactions, 5μL of PCR amplicon was mixed with 3μL nuclease-free water (Thermo Fisher Scientific) and 2μL ExoSAP (Exonuclease I and rSAP, New England Biolabs), then incubated at 37°C for 30 minutes to degrade primers and dephosphorylate dNTPs, and enzymes were then inactivated via incubation at 80°C for 15 minutes at MIT. At SIAT, 5uL of PCR amplicon was mixed with 2uL ExoSAP-IT reagent (Thermo Fisher Scientific), then incubated at 37°C for 15 minutes to degrade primers and dephosphorylate dNTPs, and enzymes were inactivated via incubation at 80°C for 15 minutes. 5μL of 5μM sequencing primer was then added to each mixture and samples were submitted to Genewiz for Sanger sequencing. Sequence files (.ab1) were analyzed using SnapGene. To quantify the frequencies of intended mutations, ab1 files were analyzed using ICE software (Synthego, https://ice.synthego.com/#/).

#### in silico FEN1 Expression Analysis

The data used for the *FEN1* expression analyses described in this manuscript were obtained from the Genotype-Tissue Expression (GTEx) Portal (Release V6) (https://www.gtexportal.org/home/) and Human Protein Atlas (http://www.proteinatlas.org).

#### Statistical Analysis

Statistical analysis was performed with Prism 7 (GraphPad).

