## Supplementary Information for "Prime editing primarily induces undesired outcomes in mice"

***Prime Editing with T7 in vitro Transcribed pegRNAs and Nicking sgRNAs*:** We performed in vitro fertilization of mouse embryos, injected mRNA encoding the PE2 enzyme along with T7 in vitro transcribed (T7-IVT) pegRNA and nicking-sgRNA, and cultured the embryos until the blastocyst stage before DNA isolation and genotyping (Figure S1). For deep analysis of the resulting genotypes, we performed both bulk Sanger sequencing of PCR products from total blastocyst DNA and also sequenced cloned PCR products to assess individual alleles (see STAR methods). In our initial trial, we did not observe clear modification of any locus (data not shown), consistent with the very low modification frequencies in mice recently reported by the others (Liu et al., 2020). We hypothesized that the standard T7-pegRNA template amplification by PCR (Aida et al., 2015; Wang et al., 2013) might not be suitable for pegRNA production since the T7-spacer forward primer can bind to both the bona fide spacer and primer-binding site (PBS) in the 3' end of pegRNA, making correct PCR amplification less efficient. Therefore, we cloned T7 sequences to the pegRNA expression plasmids, linearized, and ran T7-IVT to produce pegRNAs (see STAR methods). By using these IVT pegRNAs derived from T7-plasmids, we observed low-to-moderate modifications of the Dnmt1 and Chd2 loci, but not Tyr (Figure S2A). Approximately 10% and 30% of PE3-injected embryos were positive for the intended edit with approximately 40% and 20% median allele frequencies in Dnmt1 and Chd2, respectively (Figures S2B and S2C). Remarkably, half of the modified Chd2 embryos had undesired indels (Figure S2D).

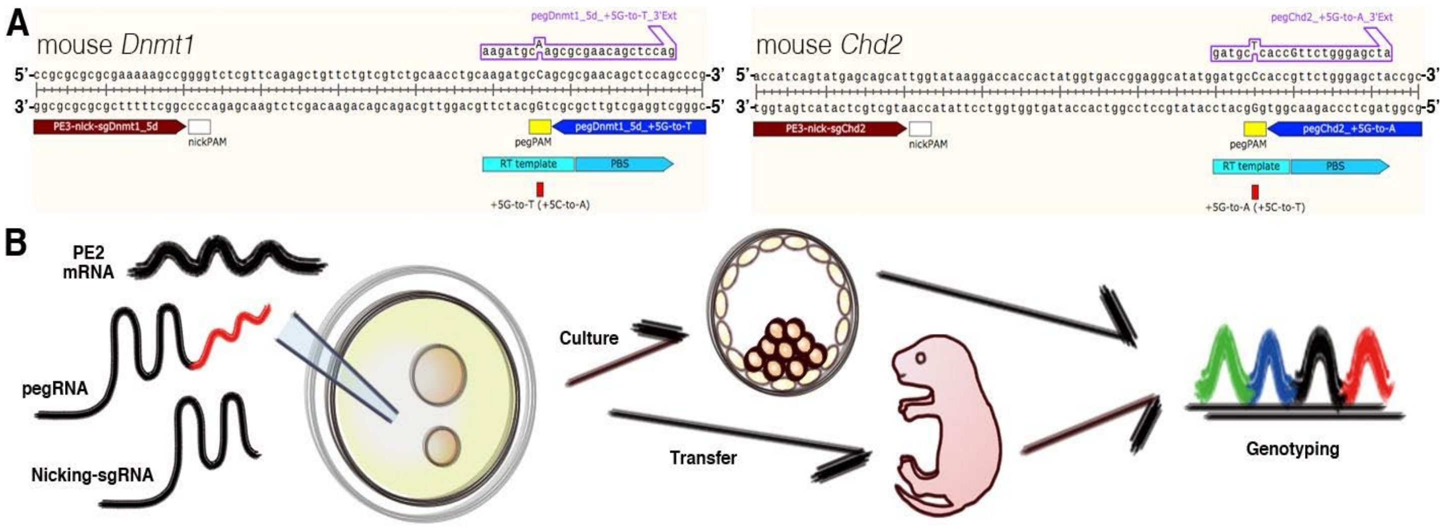


**Figure S1. Prime editing strategy in mouse embryos, Related to Figure 1.** (**A**) Schematic of *Dnmt1* and *Chd2* loci depicting the locations of pegRNA and sgRNA protospacers, as well as the intended prime edits to be installed and their corresponding reverse transcription (RT) templates. PBS: primer-binding site. (**B**) Cytoplasmic injection strategy for generating and genotyping prime edited embryos and pups.

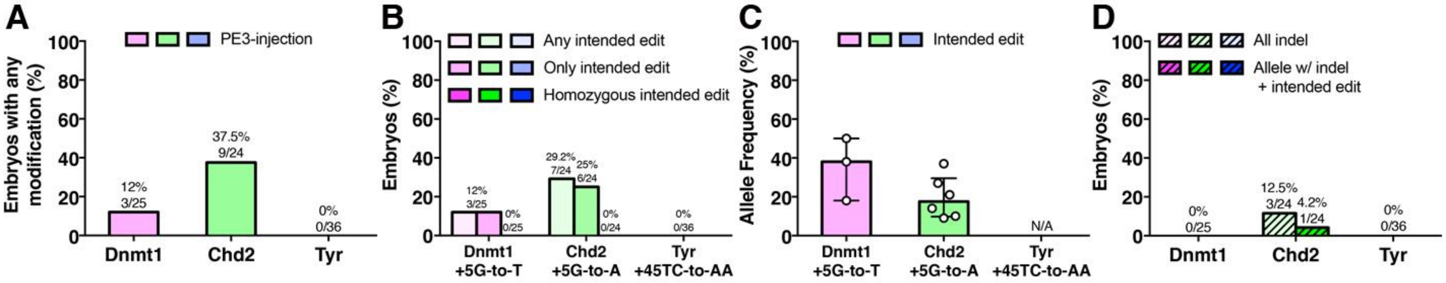


**Figure S2. Prime editing by T7-IVT pegRNA and nicking-sgRNA in mouse embryos, Related to Figure 1.** (**A,B**) The percentage of embryos carrying (A) any modification of the target site or (B) the intended edit at the target site. (**C**) Allele frequencies of the intended edits (see **STAR methods**). (**D**) Percentage of embryos with either indels or individual alleles carrying both an indel and the intended edit. Values shown are median ± interquartile range. N/A: not applicable.

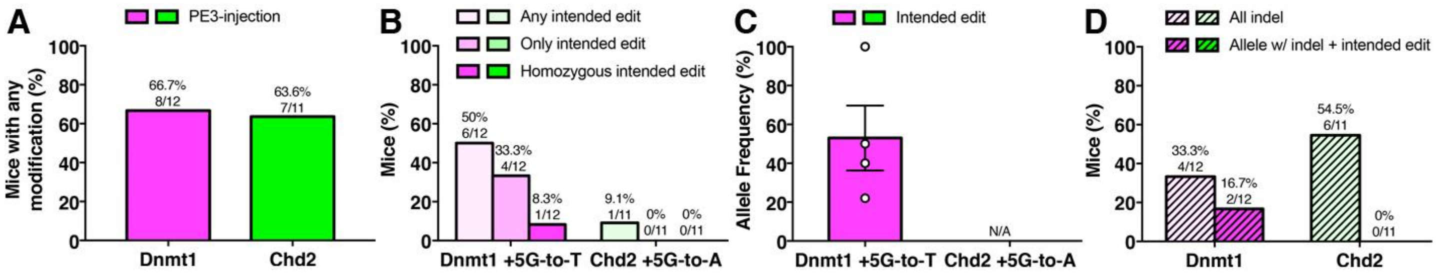


**Figure S3. Prime Editing by PE3 in newborn mice, Related to Figure 1.** (**A,B**) The percentage of mice carrying (A) any modification of the target site or (B) the intended edit at the target site. (**C**) Allele frequencies of the intended edits (see **STAR methods**). (**D**) Percentage of mice with either indels or individual alleles carrying both an indel and the intended edit. Values shown are median ± interquartile range. N/A: not applicable.

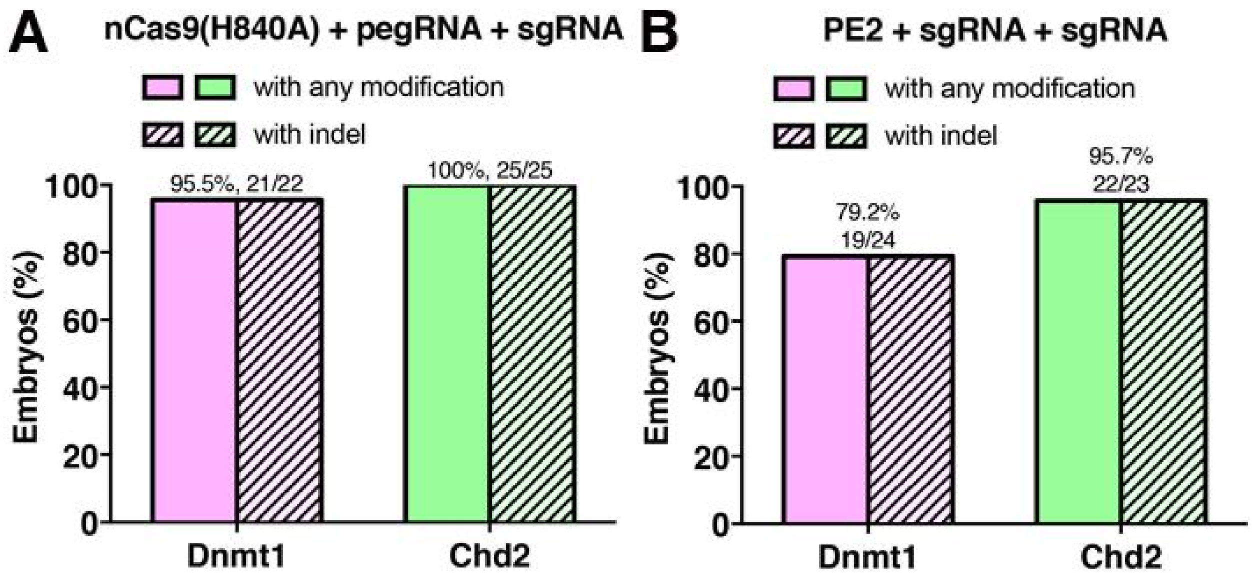


**Figure S4. Double nicking induces indels, Related to Figure 1.** (**A**,**B**) The percentage of embryos carrying any modification of the target site or indel by double nicking using (A) PE2 lacking RT domain (equivalent to Cas9 nickase (H840A)), pegRNA and nicking-sgRNA or (B) PE2, pegRNA lacking RTT and PBS (equivalent to sgRNA) and nicking-sgRNA.

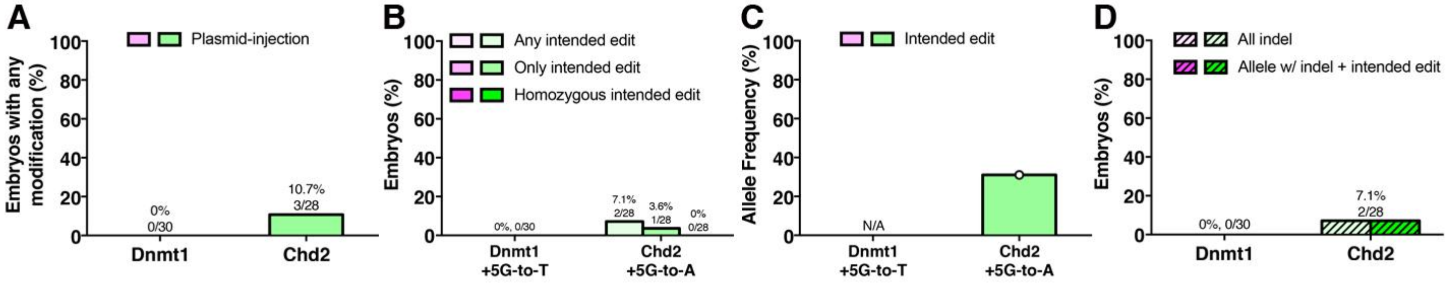


**Figure S5. Prime editing by DNA injection in mouse embryos, Related to Figure 1.** (**A,B**) The percentage of embryos carrying (A) any modification of the target site or (B) the intended edit at the target site. (**C**) Allele frequency of the intended edits (see **STAR methods**). (**D**) Percentage of embryos with either indels or individual alleles carrying both an indel and the intended edit. Value shown is median. N/A: not applicable.

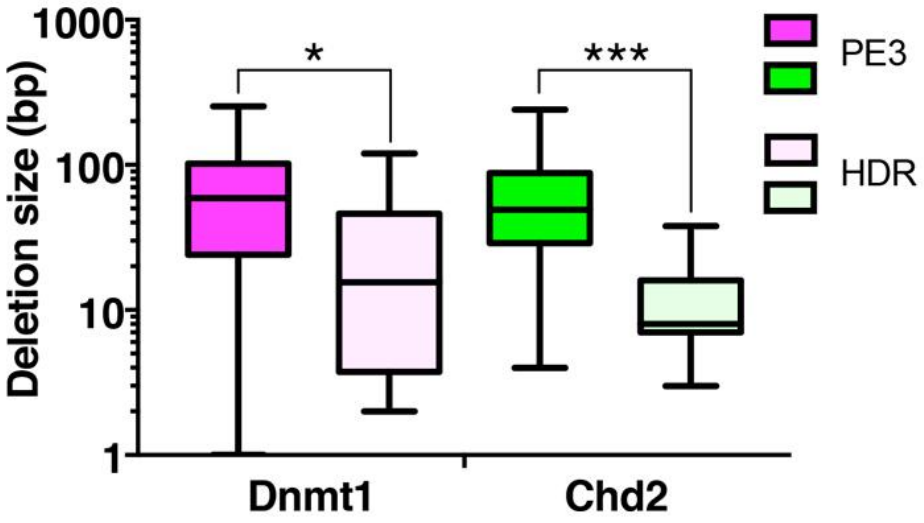


**Figure S6. Comparison of deletion sizes between PE3 and HDR, Related to Figures 1 and 4.** Comparison of cumulative distributions of observed deletion sizes with either PE3 (bright colors) or HDR (dull colors) editing strategies (Kolmogorov-Smirnov test; * - $p<0.05$; *** - $p<0.001$). Data from Figures 1H and 4I.

| pegRNA | Sequence |
|---|---|
| pegDnmt1_5d_5GtoT | mC*mG*mG*GCUGGAGCUGUUCGCGCGUUUUAGAGCUAGAAAUAGCAAGUUAAAAUAAGGCUAGUCCGUUAU<br>CAACUUGAAAAAGUGGCACCGAGUCGGUGCAAGAUGCAAGCGCGAACAGCUCCAG*mU*mU*mU |
| pegChd2_5GtoA | mG*mC*mG*GUAGCUCCCAGAACGGUGUUUUAGAGCUAGAAAUAGCAAGUUAAAAUAAGGCUAGUCCGUUAU<br>CAACUUGAAAAAGUGGCACCGAGUCGGUGCGAUGCUCACCGUUCUGGGAGCUA*mU*mU*mU |
| pegRnf2_1CG | mA*mG*mG*AUGUAUUAUAUUACCUGGUUUUAGAGCUAGAAAUAGCAAGUUAAAAUAAGGCUAGUCCGUUAU<br>CAACUUGAAAAAGUGGCACCGAGUCGGUGCAACGAACACCUCACGUAAUAUAAUACAUC*mU*mU*mU |
| pegRnf2_1GTAins | mA*mG*mG*AUGUAUUAUAUUACCUGGUUUUAGAGCUAGAAAUAGCAAGUUAAAAUAAGGCUAGUCCGUUAU<br>CAACUUGAAAAAGUGGCACCGAGUCGGUGCAACGAACACCUCAGUACGUAAUAUAAUACAUC*mU*mU*mU |
| pegRnf2_3-5GAGdel | mA*mG*mG*AUGUAUUAUAUUACCUGGUUUUAGAGCUAGAAAUAGCAAGUUAAAAUAAGGCUAGUCCGUUAU<br>CAACUUGAAAAAGUGGCACCGAGUCGGUGCAACGAACACAGGUAAUAUAAUACAUC*mU*mU*mU |
| pegTyr_45TCtoAA | mG*mA*mA*GUUGCCUGAGCACUGGCGUUUUAGAGCUAGAAAUAGCAAGUUAAAAUAAGGCUAGUCCGUUAU<br>CAACUUGAAAAAGUGGCACCGAGUCGGUGCAAUAGGACAAGCCAGUGCUCAGGCAA*mU*mU*mU |
| pegActb_1GAATTC | mC*mA*mU*UAUGAGUCCUUAAGUGAGUUUUAGAGCUAGAAAUAGCAAGUUAAAAUAAGGCUAGUCCGUUAU<br>CAACUUGAAAAAGUGGCACCGAGUCGGUGCUGCAUUUGCCUUCAGAAUUCCUUAAGGACUCAU*mU*mU*mU |
| pegCol12a1_2AtoC | mU*mG*mA*CUUCCAUGGUUCCACAAGUUUUAGAGCUAGAAAUAGCAAGUUAAAAUAAGGCUAGUCCGUUAU<br>CAACUUGAAAAAGUGGCACCGAGUCGGUGCAAUGGACCCAUGGUGGAACCAUGGAA*mU*mU*mU |
| pegDnmt1_5d_3b_6GtoC | mC*mG*mG*GCUGGAGCUGUUCGCGCGUUUUAGAGCUAGAAAUAGCAAGUUAAAAUAAGGCUAGUCCGUUAU<br>CAACUUGAAAAAGUGGCACCGAGUCGGUGCAAGAUGGCAGCGCGAACAGCUCCAG*mU*mU*mU |
| pegChd2_3b_6GtoC | mG*mC*mG*GUAGCUCCCAGAACGGUGUUUUAGAGCUAGAAAUAGCAAGUUAAAAUAAGGCUAGUCCGUUAU<br>CAACUUGAAAAAGUGGCACCGAGUCGGUGCGAUGGCCACCGUUCUGGGAGCUA*mU*mU*mU |

| sgRNA | Sequence |
|---|---|
| sgDnmt1_5d_Nick | mC*mC*mG*CGCGCGCGAAAAAGCCGGUUUUAGAGCUAGAAAUAGCAAGUUAAAAUAAGGCUAGUCCGUUAU<br>CAACUUGAAAAAGUGGCACCGAGUCGGUGCU*mU*mU*mU |
| sgDnmt1_5d_5GtoT_DSB | mC*mG*mG*GCUGGAGCUGUUCGCGCGUUUUAGAGCUAGAAAUAGCAAGUUAAAAUAAGGCUAGUCCGUUAU<br>CAACUUGAAAAAGUGGCACCGAGUCGGUGCU*mU*mU*mU |
| sgChd2_Nick | mA*mC*mC*AUCAGUAUGAGCAGCAUGUUUUAGAGCUAGAAAUAGCAAGUUAAAAUAAGGCUAGUCCGUUAU<br>CAACUUGAAAAAGUGGCACCGAGUCGGUGCUU*U*U* |
| sgChd2_5GtoA_DSB | mG*mC*mG*GUAGCUCCCAGAACGGUGUUUUAGAGCUAGAAAUAGCAAGUUAAAAUAAGGCUAGUCCGUUAU<br>CAACUUGAAAAAGUGGCACCGAGUCGGUGCU*mU*mU*mU |
| sgRnf2_Nick41 | mU*mC*mA*ACCAUUAAGCAAAACAUGUUUUAGAGCUAGAAAUAGCAAGUUAAAAUAAGGCUAGUCCGUUAU<br>CAACUUGAAAAAGUGGCACCGAGUCGGUGCU*mU*mU*mU |
| sgRnf2_Nick67 | mU*mC*mU*CAGGCUGUGCAGACAAAGUUUUAGAGCUAGAAAUAGCAAGUUAAAAUAAGGCUAGUCCGUUAU<br>CAACUUGAAAAAGUGGCACCGAGUCGGUGCU*mU*mU*mU |
| sgTyr_Nick | mG*mG*mA*CCUCAGUUCCCCUUCAAGUUUUAGAGCUAGAAAUAGCAAGUUAAAAUAAGGCUAGUCCGUUAU<br>CAACUUGAAAAAGUGGCACCGAGUCGGUGCU*mU*mU*mU |
| sgActb_Nick | mA*mG*mA*UCCAGUGCUCUUUAGCAGUUUUAGAGCUAGAAAUAGCAAGUUAAAAUAAGGCUAGUCCGUUAU<br>CAACUUGAAAAAGUGGCACCGAGUCGGUGCU*mU*mU*mU |
| sgCol12a1_Nick | mG*mC*mC*UGAGCAGGCCACGAACAGUUUUAGAGCUAGAAAUAGCAAGUUAAAAUAAGGCUAGUCCGUUAU<br>CAACUUGAAAAAGUGGCACCGAGUCGGUGCUU*U*U* |
| sgDnmt1_3b_6GtoC_Nick | mG*mU*mC*GUCUGCAACCUGCAAGAGUUUUAGAGCUAGAAAUAGCAAGUUAAAAUAAGGCUAGUCCGUUAU<br>CAACUUGAAAAAGUGGCACCGAGUCGGUGCU*mU*mU*mU |
| sgChd2_3b_6GtoC_Nick | mG*mG*mU*GACCGGAGGCAUAUGGAGUUUUAGAGCUAGAAAUAGCAAGUUAAAAUAAGGCUAGUCCGUUAU<br>CAACUUGAAAAAGUGGCACCGAGUCGGUGCU*mU*mU*mU |

**Table S1. Sequences of chemically synthesized pegRNAs and sgRNAs, Related to STAR Methods.** Red: protospacers, pink: intended mutation, blue: reverse-transcription template, green: primer-binding site, *: 2'-O-methyl analogs and 3'-phosphorothioate internucleotide linkages.

| pegRNA | Sequence |
|---|---|
| T7-pegDnmt1_5d_5GtoTspacerS | CACCTAATACGACTCACTATAGCGGGCTGGAGCTGTTCGCGCGTTTT |
| T7-pegDnmt1_5d_5GtoTspacerAs | CTCTAAAACGCGCGAACAGCTCCAGCCCGCTATAGTGAGTCGTATTA |
| T7-pegChd2_5GtoAspacerS | CACCTAATACGACTCACTATAGGCGGTAGCTCCCAGAACGGTGTTTT |
| T7-pegChd2_5GtoAspacerAs | CTCTAAAACACCGTTCTGGGAGCTACCGCCTATAGTGAGTCGTATTA |
| T7-pegTyr_45TCtoAAspacerS | CACCTAATACGACTCACTATAGGAAGTTGCCTGAGCACTGGCGTTTT |
| T7-pegTyr_45TCtoAAspacerAs | CTCTAAAACGCCAGTGCTCAGGCAACTTCCTATAGTGAGTCGTATTA |
| pegRNA scaffold-S | Phos-AGAGCTAGAAATAGCAAGTTAAAATAAGGCTAGTCCGTTATCAACTTGAAAAAGTGGCACCGAGTCG |
| pegRNA scaffold-As | Phos-GCACCGACTCGGTGCCACTTTTTCAAGTTGATAACGGACTAGCCTTATTTTAACTTGCTATTTCTAG |
| pegDnmt1_5d_5GtoT3extS | GTGCAAGATGCAAGCGCGAACAGCTCCAG |
| pegDnmt1_5d_5GtoT3extAs | AAAACTGGAGCTGTTCGCGCTTGCATCTT |
| pegChd2_5GtoA3extS | GTGCGATGCTCACCGTTCTGGGAGCTA |
| pegChd2_5GtoA3extAs | AAAATAGCTCCCAGAACGGTGAGCATC |
| pegTyr_45TCtoAA3extS | GTGCAATAGGACAAGCCAGTGCTCAGGCAA |
| pegTyr_45TCtoAA3extAs | AAAATTGCCTGAGCACTGGCTTGTCCTATT |

| sgRNA | Sequence |
|---|---|
| Nick-sgDnmt1_5d-S | CACCGGCCGCGCGCGCGAAAAAGCCG |
| Nick-sgDnmt1_5d-As | AAACCGGCTTTTTCGCGCGCGCGGCC |
| T7-Nick-sgDNMT1_5d-F | TAATACGACTCACTATAGGGCCGCGCGCGCGAAAAAGCCG |
| Nick-sgChd2-S | CACCGACCATCAGTATGAGCAGCAT |
| Nick-sgChd2-As | AAACATGCTGCTCATACTGATGGTC |
| T7-Nick-sgChd2-F | TAATACGACTCACTATAGGGACCATCAGTATGAGCAGCAT |
| T7-Nick-sgTyr-sgF | TAATACGACTCACTATAGGGACCTCAGTTCCCCTTCAAGTTTTAGAGCTAGAAATAGC |
| tracrRNA-R | AAAAGCACCGACTCGGTGCC |

| Cas9 | Sequence |
|---|---|
| PE2-RTremove-S | TGTCTCAGCTGGGAGGTGACCCCAAGAAGAAGAGGAAAGT |
| PE2-RTremove-As | ACTTTCCTCTTCTTCTTGGGGTCACCTCCCAGCTGAGACA |
| T7-Cas9-F | TAATACGACTCACTATAGGGAGAGCCG |
| SV40NLS-Cas9-R | TTAGACTTTCCTCTTCTTCTTGGGGTCACCTCCCAGCTGAGAC |

| HDR donor | Sequence |
|---|---|
| ssDnmt1_5d_5GtoT_S_HDR | CGCGCGAAAAAGCCGGGGTCTCGTTCAGAGCTGTTCTGTCGTCTGCAACCTGCAAGATGCAAGCGCGAACAG CTCCAGCCCGAGTGCCTGCGCTTGCCTCCCCGGCAGGCTCGCTCCCGGA |
| ssChd2_RH_S_HDR | TCAGTATGAGCAGCATTGGTATAAGGACCACCACTATGGTGACCGGAGGCATATGGATGCTCACCATTCTGG GAGCTACCGCCCTAACAACATGTCCAGAAAGAGGCCGTATGAGCAGTACAACAG |

**Table S2. Sequences of oligos for *in vitro* transcription and HDR donor, Related to STAR Methods.**
Pink: intended mutation

| Primer | Sequence |
|---|---|
| Dnmt1-F1 | GCCTCTGCCAGACTCAAAAG |
| Dnmt1-R1 | CCTCAAGCTCCCAGTCAATG |
| Dnmt1-F2 | ATGTGAGTGCAATGCCCATA |
| Dnmt1-R2 | ACCCGGAGATACCCCCAATA |
| Chd2-F1 | GGGGTGAAATTACCAAAAGC |
| Chd2-R1 | TGCAGATTAGGGCAAGCAAG |
| Chd2-F2 | GACAATGCATCCTCCTCTCC |
| Chd2-R2 | TCTTGCCTTCAAAGTTGGTG |
| Tyr-F1 | TCAATTTAGTTACCTCACTATGGGC |
| Tyr-R1 | CAAGTACTCATCTGTGCAAATGTC |
| Tyr-F2 | TTTGGCCATAGGTGCCTG |
| Tyr-R2 | GAGCCTGTGCCTCCTCTAAG |
| Rnf2-F1 | AAGCCTTTGGATGAAATCAGAC |
| Rnf2-R1 | AGATGCTGAAGACTGGAGGG |
| Rnf2-F2 | CAACAGTGTCACTCGCCC |
| Rnf2-R2 | TTGGACACCAGTTTGAGGAATAC |
| Actb-F1 | CTGACAGCAGGAAGGTGTGA |
| Actb-R1 | TCTTTTGCAGCAAGGTCTCA |
| Acbt-F2 | TGCAGAGAACACTGGTTGGT |
| Actb-R2 | GGCTGGCCTCAAACTCAGTA |
| Col12a1-F1 | TCAAATTACCCTGGGACCAC |
| Col12a1-R1 | AAGGTCACAGTCCTGACCCA |
| Col12a1-F2 | CCCAGCTGGAAGACACATTT |
| Col12a1-R2 | CAGACCAAGCAACAGAAGCA |
| M13-20 | GTAAAACGACGGCCAGT |
| M4668 | CAGGAAACAGCTATGACCATGAT |

**Table S3. Sequences of PCR primers, Related to STAR Methods.**
